# spillR: Spillover Compensation in Mass Cytometry Data

**DOI:** 10.1101/2023.10.04.560870

**Authors:** Marco Guazzini, Alexander G. Reisach, Sebastian Weichwald, Christof Seiler

**Affiliations:** Department of Advanced Computing Sciences, Maastricht University, The Netherlands; Université Paris Cité, CNRS, MAP5, F-75006 Paris, France; Department of Mathematical Sciences, University of Copenhagen, Denmark; Mathematics Centre Maastricht, Maastricht University, The Netherlands; Center of Experimental Rheumatology, Department of Rheumatology, University Hospital Zurich, University of Zurich, Switzerland

## Abstract

Channel interference in mass cytometry can cause spillover and may result in miscounting of protein markers. Chevrier *et al*. (2018) introduce an experimental and computational procedure to estimate and compensate for spillover implemented in their R package `CATALYST`. They assume spillover can be described by a spillover matrix that encodes the ratio between unstained and stained channels. They estimate the spillover matrix from experiments with beads. We propose to skip the matrix estimation step and work directly with the full bead distributions. We develop a nonparametric finite mixture model, and use the mixture components to estimate the probability of spillover. Spillover correction is often a pre-processing step followed by downstream analyses, choosing a flexible model reduces the chance of introducing biases that can propagate downstream. We implement our method in an R package `spillR` using expectation-maximization to fit the mixture model. We test our method on synthetic and real data from `CATALYST`. We find that our method compensates low counts accurately, does not introduce negative counts, avoids overcompensating high counts, and preserves correlations between markers that may be biologically meaningful.

## 1 Introduction

Mass cytometry makes it possible to count a large number of proteins simultaneously on individual cells (Bandura et al., 2009; Bendall et al., 2011). Although mass cytometry has less spillover—measurements from one channel overlap less with those of another—than flow cytometry (Bagwell and Adams, 1993; Novo et al., 2013), spillover is still a problem and affects downstream analyses such as differential testing (Weber *et al*., 2019; Seiler et al., 2021) or dimensionality reduction (McCarthy *et al*., 2017). Reducing spillover by careful design of experiment is possible (Takahashi *et al*., 2017), but a purely experimental approach may be neither sufficient nor efficient (Lun *et al*., 2017).

Chevrier *et al*. (2018) propose a method for addressing spillover by conducting an experiment on beads. This experiment measures spillover by staining each bead with a single type of antibody. The slope of the regression line between target antibody and non-target antibodies represents the spillover proportion between channels. Miao *et al*. (2021) attempt to solve spillover by fitting a mixture model. Our contribution combines the solutions of Chevrier *et al*. (2018) and Miao *et al*. (2021). We still require a bead experiment, as in Chevrier *et al*. (2018), but estimate spillover leveraging a statistical model, as in Miao *et al*. (2021). Both previous versions rely on an estimate for the spillover matrix. The spillover matrix encodes the pairwise spillover proportion between channels. We avoid estimating a spillover matrix and instead model spillover by fitting a mixture model to the observed counts. Our main new assumption is that the spillover distribution—not just the spillover proportion—from the bead experiment carries over to the biological experiment. In other words, we transfer the spillover distribution to the real experiment instead of just the spillover proportion encoded in the spillover matrix. The main difference between our method and the one by Chevrier *et al*. (2018) is the average count after correcting for spillover. Our method will increase the average count after correction. By contrast, the method by Chevrier *et al*. (2018) will shrink all the counts towards zero. As a consequence, the average count will be lower after correction.

In Section 2, we present our mixture model and link it to calculating spillover probabilities for specific count values. Our estimation procedure is based on an EM algorithm and logistic regression, and implemented in our new R package `spillR`^1^. In Section 3, we conduct experiments on simulated and real data obtained from the `CATALYST R` package (Chevrier *et al*., 2018). Section 4 discusses our synthetic experiments and relates our findings to `CATALYST`.

## 2 Methods

### 2.1 Example

Figure 1 illustrates our procedure using a dataset from the `CATALYST` package as an example. There are four markers, HLA-DR (Yb171Di), HLA-ABC (Yb172Di), CD8 (Yb174Di), and CD45 (Yb176Di), that spill over into the target marker, CD3 (Yb173Di). The markers have two names: the first name is the protein name and the second name in brackets is the conjugated metal. There are bead experiments for each of the spillover markers.

**Figure 1.**
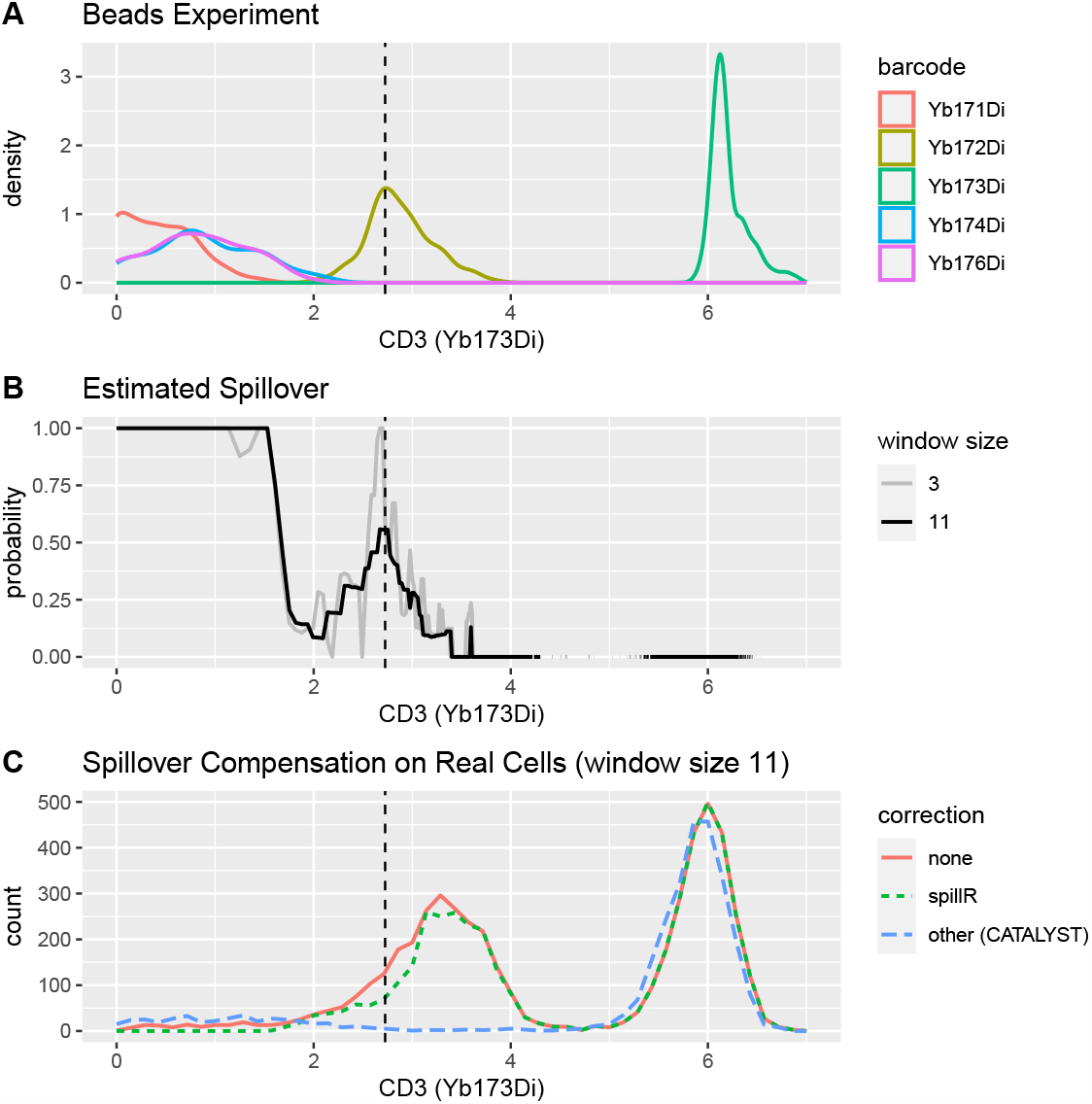
A: Gaussian kernel density plot of target and spillover markers. B: The solid black curve represents the spillover probability estimated with our method with smoothing parameter *k* = 11. The grey curve is the estimate with *k* = 3. Larger values yield smoother spillover probability curves. C: Frequency polygons of marker CD3 (Yb173Di) comparing no correction, our method, and another method. Zero counts are not plotted. See Table 1 for the zero counts and means. Counts are arcsinh transformed with cofactor of five (Bendall *et al*., 2011). The vertical dashed line helps to interpret the spillover correction when CD3 (Yb173Di) is 2.725. This is an interesting point as it balances between spillover counts from Yb172Di measured in beads (panel A) and the counts measured on real cells (panel C). The spillover probability is around 0.5. This means we will set about half of the counts to `NA`’s at this point. After correction the read curve (panel C) will be adjusted downwards to the green curve (panel C) with our method `spillR`.

Panel A depicts the marker distributions from the beads experiment. We see that for this marker the bead experiments are high-quality as the target marker Yb173Di is concentrated around six, similarly to the experiment with real cells. This suggests that the spillover marker values can be transferred to the real experiments. Marker Yb172Di shows large spillover into Yb173Di, and suggests that the left tail of the first mode of the distribution may be attributed to that marker. The other spillover markers have low counts, making it justifiable to set some or all the low counts to zero.

Panel B has a solid black and gray curves representing our spillover probability estimates. With the smoothing parameter *k* = 11, we can see that the probability of spillover goes up to around 0.5. In that case, our correction step assigns around 50% of cells to spillover, and keeps the other 50% at the current value. We add a black dashed line to all plots at the position 2.725 to illustrate this point. Low counts have spillover probability of one, which means that our procedure assigns them to spillover and masks them from the sample by setting them to `NA` values. Counts above four stem from spillover with probability zero (and from the actual target with probability one), which means that our procedure keeps them at their raw uncorrected value. The estimated curve with smoothing parameter *k* = 3 is more irregular, suggesting that these fluctuations may be driven by noise. Users can control the smoothing parameter *k* to choose the desired bias-variance tradeoff.

Panel C displays the distribution of our target marker, CD3 (Yb173Di), before and after spillover correction. Our compensation method, `spillR`, masks most markers in the low counts up to two, about 25% to 50% in the medium counts range between two and four, and keeps counts above four. Larger counts are not affected by the correction, as can be seen by the overlap of both curves. In contrast, the other method, `CATALYST`, shifts large counts to the left, and shifts medium counts to low counts or zero counts. Zero counts (or equivalently `NA` counts for `spillR`) and mean counts are shown in Table 1.

**Table 1:** Additional summaries of the data underlying Figure 1C.

| correction | zeros or NA's | mean |
| --- | --- | --- |
| none | 31 | 4.45 |
| **spillR** | 511 | 4.71 |
| other (CATALYST) | 2162 | 2.94 |

### 2.2 Definition of Spillover Probability and Assumptions

We observe a count *Y*_*i*_ of a target marker in cell *i*. We model the observed *Y*_*i*_ as a finite mixture (McLachlan *et al*., 2019) of unobserved true marker counts *Y*_*i*_|*Z*_*i*_ = 1 and spillover marker counts *Y*_*i*_|*Z*_*i*_ = 2, …, *Y*_*i*_|*Z*_*i*_ = *K* with mixing probabilities *π*_*k*_ = *P* (*Z*_*i*_ = *k*) for *k* = 1, …, *K*,

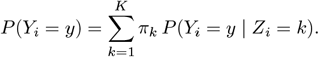

The first mixing probability is the proportion of true signal in the observed counts. The other *K −*1 mixing probabilities are the proportions of spillover. The total sum of mixing probabilities equals one, ∑_*k*_ *π*_*k*_ = 1. The total number of markers in mass cytometry panels is between 30 and 40 (Bendall *et al*., 2011), but only a small subset of three to four markers spill over into the target marker (Chevrier *et al*., 2018). So, typically *K* = 1 + 3 or *K* = 1 + 4.

Experimentally, we only measure a sample from the distribution of *Y*_*i*_. The probabilities *π*_*k*_ and true distributions *P* (*Y*_*i*_ = *y*|*Z*_*i*_ = *k*) are unobserved, and we need to estimate them from data. In many applications, the mixture components are in a parametric family, for example, the negative binomial distribution. As spillover correction is a pre-processing step followed by downstream analyses, choosing the wrong model can introduce biases in the next analysis step. To mitigate such biases, we propose to fit nonparametric mixture components. We make two assumptions that render the components and mixture probabilities identifiable:

- (A1) Spillover distributions are the same in bead and real experiments. The distribution of *Y*_*i*_|*Z*_*i*_ = *k* for all *k >* 1 is the same in beads and real cells. This assumption allows us to learn the spillover distributions of *Y*_*i*_ |*Z*_*i*_ = *k* for all *k >* 1 from experiments with beads, and transfer them to the experiment with real cells. This assumption relies on high-quality single stained bead experiments that measure spillover in the same range as the target biological experiment. In other words, a high-quality bead experiment for our method works best if the distribution of bead cells is similar to the distribution of real cells.
- (A2) For each cell *i*, the observed count *Y*_*i*_ can only be due to one distribution.

This assumption is already implied by the statement of the mixture model. It allows us to calculate the spillover probability for a given count *Y*_*i*_ = *y* from the posterior probability that it arises through spillover from markers *k >* 1,

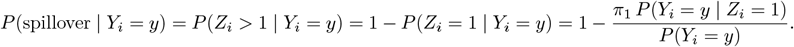

To parse this calculation, recall that in mixture models the *π*_1_ is the prior probability, *P* (*Y*_*i*_ = *y*|*Z*_*i*_ = 1) is the conditional probability given the mixture component, and the denominator *P* (*Y*_*i*_ = *y*) is the marginal distribution. Applying Bayes rule leads to the posterior probability.

### 2.3 Estimation of Spillover Probability

We propose a two step procedure for estimating the spillover probability. In step 1, we estimate mixture components and mixture probabilities. We refine these estimates using the EM algorithm (Dempster *et al*., 1977). In step 2, we use these probability estimates to assign counts to spillover or signal.

We denote the *n×K* count matrix as **Y** = (*y*_*ik*_) with real cells in the first column and beads in columns two and higher. To simplify mathematical notation but without loss of generality, we assume that the number of cells from real and bead experiments have the same *n*. In practice, the number of cells from bead experiments is much smaller than from real experiments. The *k*th column of **Y** contains marker counts for the *k*th spillover marker, which represents the empirical spillover distribution of marker *k* into the target marker, that is, the marker in the first column of **Y**.

We use the empirical weighted cumulative distribution function (CDF) with non-negative weights *w*_*i*_ at each data point to model the marker distributions *F* for a marker *k*,

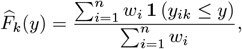

where **1**(*·*) is the indicator function.

#### 2.3.1 EM Algorithm

- Initialization: For the mixture probability vector, we assign probability 0.9 to the the target marker and divide the probability 0.1 among the spillover markers,

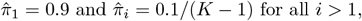

and evaluate the *k*th mixture component using its empirical CDF with equal weights for all data points,

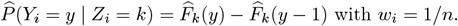

We smooth the probability mass function (PMF), 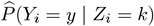, using running medians with a fixed window size as implemented in R function `runmed`. The procedure is not sensitive to the choice of the initial mixture probability vector. Other settings are possible, e.g., setting all probabilities to the same value.
- E-step: We evaluate the posterior probability of a count *y* belonging to component *k* (that is, originating from marker *k*),

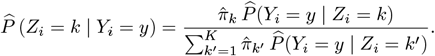
- M-step: We estimate the new mixture probability vector from posterior probabilities,

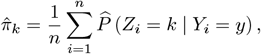

and estimate the new target marker distribution using its empirical CDF with weights set to posterior probabilities,

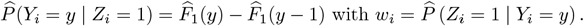

As before, we smooth the PMF, 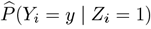, using running medians with the same window size. We keep the bead distributions, 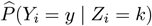 for all *k >* 1, fixed at their initial value.

To refine our estimates, we iterate over the E and M-steps until estimates stabilize. We stop iterating when 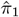 changes less than 10^−5^ from the previous iteration. The final output is the spillover probability curve with estimates at discrete points in the support of *Y*_*i*_,

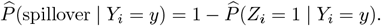

We rely on assumption (A1) to justify updating only the distribution of the target marker. We rely on assumption (A2) to justify calculating the spillover probability from the mixture model. We refer to Appendix A for a step-by-step example of our EM algorithm.

#### 2.3.2 Spillover Decision

To perform the spillover compensation, we draw from a Bernoulli distribution with the spillover probability as parameter to decide whether or not to assign a given count to spillover. We consider spillover counts as having no clear biological interpretation and mask them from our dataset while keeping all other counts. In our implementation, we choose to set spillover counts to `NA` instead of zero to avoid zero-inflated distributions.

## 3 Results

We first evaluate our new method `spillR` on simulated datasets. We probe our method to experimentally find its shortcomings. Then, we compare `spillR` to the non-negative least squares method implemented in the R package `CATALYST` on real data from the same package. All experiments and plots can be reproduced by compiling the R markdown file `spillR_paper.Rmd`^2^.

### 3.1 Simulated Data

We choose three different experiments to test spillR against different bead and real cell distributions. We explore a wide range of possible parameter settings. Figure 2 has three panels, each representing one experimental setup. The first two panels test our assumptions (A1) and (A2). The third panel tests sensitivity of `spillR` to bimodal bead distributions. For all three experiments, we model counts using a Poisson distribution with parameter *λ*. We simulate 10,000 real cells with *λ* = 100, and 1,000 beads with *λ* = 70, and spillover probability of 0.1. Beads are an independent copy of the true spillover. The other parameters and statistical dependencies are specific to each experiment. The details of the generative models are given in Appendix B. We repeat each simulation 20 times and report averages over the 20 replications. Each panel of Figure 2 has two rows of plots. The plot on the first row represents the summary of the means for each experimental setup as a function of their respective parameter *τ*. This parameter has a different meaning in each setup. To visualize the different experiments, we summarize the full distributions with the true simulated signal mean (black), the uncorrected mean (orange), and the `spillR` corrected mean (green). Plots on the second row illustrate the simulated data distributions for three selected parameters *τ* picked from the experimental setup. The yellow density curve is the observed count *Y*. The black density curve is the target cell count. The blue density curve is the spillover distribution. The goal of the experiment is to estimate the mean of the black density as accurately as possible from the yellow density curve, which represents the data *Y* that we would observe in practice. We simulate this data ourselves with the models in Appendix B.

**Figure 2.**
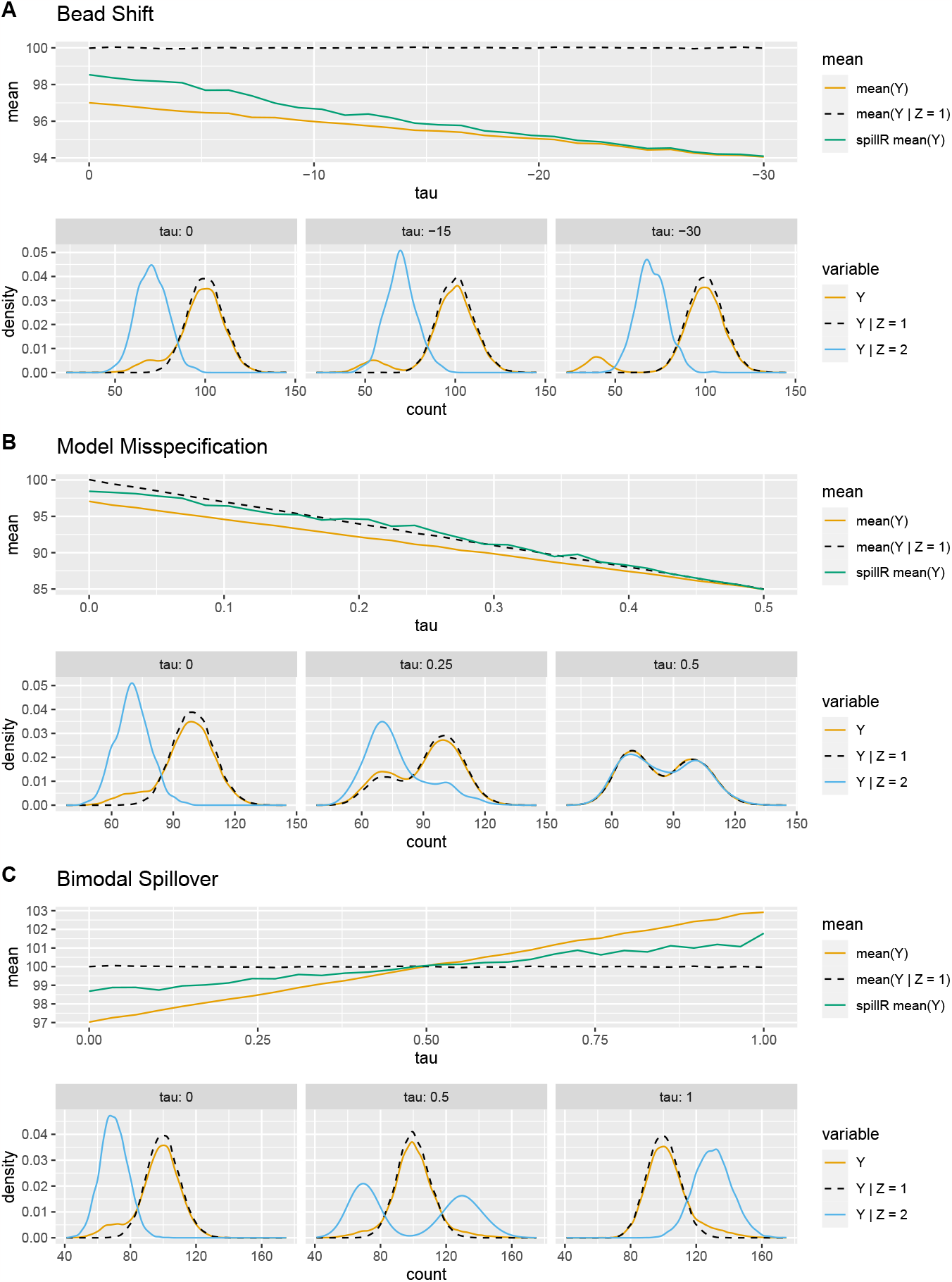
Three experiments testing our assumptions and sensitivity to bimodal bead distribution. For each experiment the top row are mean values over the entire range of the experimental setups, and the bottom row are density plots for three parameter settings to illustrate the generated distributions. *Y* is the distribution with spillover. *Y*|*Z* = 1 is the distribution without spillover. *Y*|*Z* = 2 is the spillover. mean(*Y*) is the average of the distribution with spillover. mean(*Y*|*Z* = 1) is the average count without spillover. `spillR` mean(*Y*) is the average count after correcting *Y*.

In the first experiment (panel A), we shift the measured beads spillover away from the true bead spillover to probe (A1). We test a wide range of bead shifts from no shift at *τ* = 0 to *τ* =−30. At *τ* =−30, the measured spillover (the first mode of the yellow density) is shifted away from the actual spillover (the blue density). Such low-quality beads cause both the observed and compensated mean to be below the true mean. As the beads quality improves, the compensated signal moves closer to the true mean. As we increase *τ* the first mode of the yellow density moves towards the blue density. This represents high-quality bead experiments. In all cases, even for bad beads experiments, our compensation improves the means. Our compensation also increases means, as it should, in contrast to e.g. Chevrier *et al*. (2018).

In the second experiment (panel B), we mixed target and spillover to explore the robustness of our method with respect to our second assumption (A2). One way to think about this is that the mixture is a form of model misspecification. Our mixture model is undercomplete, which means that there are more true mixture components than we observe in the beads experiment. If *τ* = 0, then assumption (A2) is correct, but for *τ* = 0.5 the assumption (A2) is maximally violated. The true mean decrease with increasing *τ*. Our compensation is closer to the true mean across the tested range. At *τ* = 0.5 all three distributions and their means are the same.

In the third experiment (panel C), we model spillover with a bimodal distribution. Here *τ* is the mixing probability of the two modes. The locations of the two spillover modes are fixed. If *τ* = 0 or *τ* = 1, then spillover is unimodal. If *τ* = 0.5, the first mode of the bimodal beads distribution is left to the signal mode, and the second mode is to the right. The corrected mean is closer to the true mean than the uncorrected mean across the test range.

### 3.2 Real Data

We compare our method to `CATALYST` on one of the example datasets in the `CATALYST` package. The dataset has an experiment with real cells and a corresponding bead experiment. The experiment on real cells has 5,000 peripheral blood mononuclear cells from healthy donors measured on 39 channels. The experiment on beads has 10,000 cells measured on 36 channels. They have single stained bead experiments. The number of beads per metal label range from 112 to 241.

We compare the two methods on the same markers as in the original `CATALYST` paper (Chevrier *et al*., 2018) in their Figure 3B. In the original experiment, they conjugated three proteins—CD3, CD8, and HLA-DR—with two different metal labels. They conjugated CD8 (first row in Figure 3) with Yb174Di (Yb is the metal and 174 is the number of neutrons of the isotope) and La139Di, and similarly for the other rows. On the horizontal axis, we plot the same markers as in the original paper, CD3 and HLA-ABC. We visualize the joint distributions using two-dimensional histograms.

**Figure 3.**
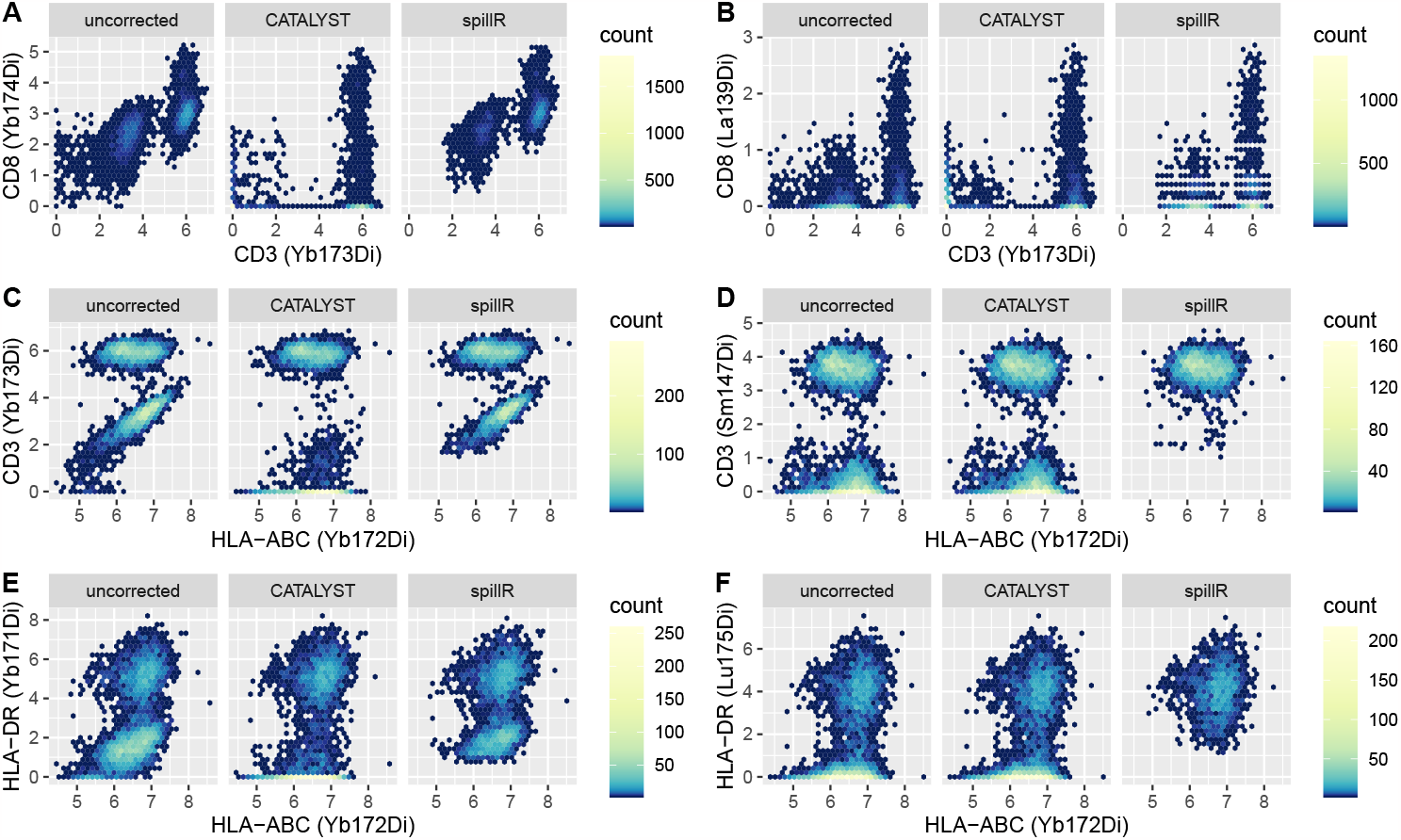
Comparison of compensation methods and uncorrected counts on real data. Counts are arcsinh transformed with cofactor of five (Bendall *et al*., 2011).

In all six panels (A–F), we observe that `spillR` compensates most strongly in the low counts. In panel C, CD3 (Yb173Di) against HLA-ABC (Yb172Di), `CATALYST` can be seen to compensate strongly in the middle range. It removes the spherical pattern that shows correlation between the two markers. `spillR` preserves this correlation structure and only masks out the lower counts of CD3 (Yb173Di). This highlights a key difference between `spillR` and `CATALYST: spillR` does not correct all counts by shrinking them, but rather removes some counts, following the idea that the distribution of the remaining counts is close to the true distribution. `CATALYST` follows another strategy by shrinking counts across the entire range.

The color code of the two-dimensional histograms indicates the absolute number of cells that fall into one hexagon bin. The uncorrected and `spillR` corrected histograms can contain different absolute numbers of cells because of how `spillR` rounds counts to integers. The raw mass cytometry data is often not count data due to proprietary post-processing of the manufacturer of the mass cytometer. That is why we convert mass cytometry data to count data, we convert the raw values to the next lower integer. The uncorrected counts do not undergo this pre-processing step. `CATALYST` does not perform this pre-processing step. This also explains the different patterns in panel B. `spillR` has horizontal stripes that correspond to non-integer values not in the support of the distribution for `spillR`.

## 4 Discussion

The experiment for (A1) shows that the mean count after `spillR` correction is closer to the true mean over a wide range of bead shifts. This indicates that our method can perform well even if the bead experiments are imperfect. If the difference between distributions of beads and real cells is large, then one option is to rerun the bead experiments to reduce this gap. The experiment for (A2) shows that our method is also robust to model misspecification. Additionally, misspecification can be addressed by adding all channels if necessary. The increase in computational cost when adding channels is relatively minor as our method scales linearly in the number of spillover markers. The experiment on bimodal bead distributions shows that the mean count after correction is still closer to the true mean even with bimodal bead distributions, and also if the spillover is actually larger than the true signal.

In our comparison to `CATALYST` on real data, we observe the effect of the two different correction strategies. `CATALYST` essentially shrinks counts towards zero by minimizing a non-negative least squares objective. It assumes that spillover is linear up to counts of 5,000. The applied shrinkage is the same for low counts (e.g., below 10) and high counts (e.g., more than 100). By contrast, `spillR` does not require linearity of the spillover, but assumes that the distribution on the beads experiment carries over to the real cells experiment. In other words, the optimal beads experiment has the same peaks as the real cells experiment.

If counts are in the spillover range (which mostly applies to low counts), they are corrected strongly and set to `NA` values. If counts are not in the spillover range, then they are left unchanged. Despite setting values to `NA`, correlations between markers are preserved. The marker correlation between HLA-ABC (Yb172Di) and CD3 (Yb173Di) illustrates this point. `CATALYST` removes the positive correlation, whereas `spillR` keeps the correlation for the higher counts. Compensation methods should try to remove spillover while keeping the biological meaningful signal for unbiased downstream analyses. Further experiments on the correlation structure between these markers are necessary to resolve the discrepancy between the two methods. This is an important point as discovering correlations between markers can lead to the discovery of new clusters or signaling networks.

Another advantage of our method is the diagnostic plot of the spillover probability curve. We can judge if the curve makes sense by comparing it to the observed count and bead distributions. Methods based on non-negative least squares are harder to diagnose as they minimize a cost function with no clear biological interpretation.

Currently, we do not take advantage of the target bead distribution in our estimation procedure. We only use spillover bead distributions. In future work, we aim to investigate ways to incorporate the target distribution into our estimation in a nonparametric manner. In our view, one of the biggest strengths of our current method is that it does not assume a specific parametric model for count data. We believe that this is crucial because spillover is just one step that precedes many downstream analysis steps, and avoiding the introduction of bias is thus our top priority.

## Acknowledgments

We thank EuroBioC2022 for awarding Marco Guazzini a travel award to present a preliminary version of `spillR` in Heidelberg. We thank Antoine Chambaz for his feedback on an earlier draft that substantially improved the paper. Alexander G. Reisach received funding from the European Union’s Horizon 2020 research and innovation program under the Marie Skłodowska-Curie grant agreement No 945332 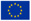.

## A EM Algorithm Example

Here we illustrate the procedure using a numerical example that includes one target and one spillover marker. We have one data matrix **Y** that contains real cell counts recorded for marker 1 (column 1) and the bead counts for marker 1 when the true marker was marker 2 (column 2). In practice, **Y** is usually a matrix with more than two columns representing multiple spillover markers. The index *i* is a specific cell in beads and real cells experiment, respectively. Let’s assume the following counts,

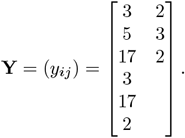

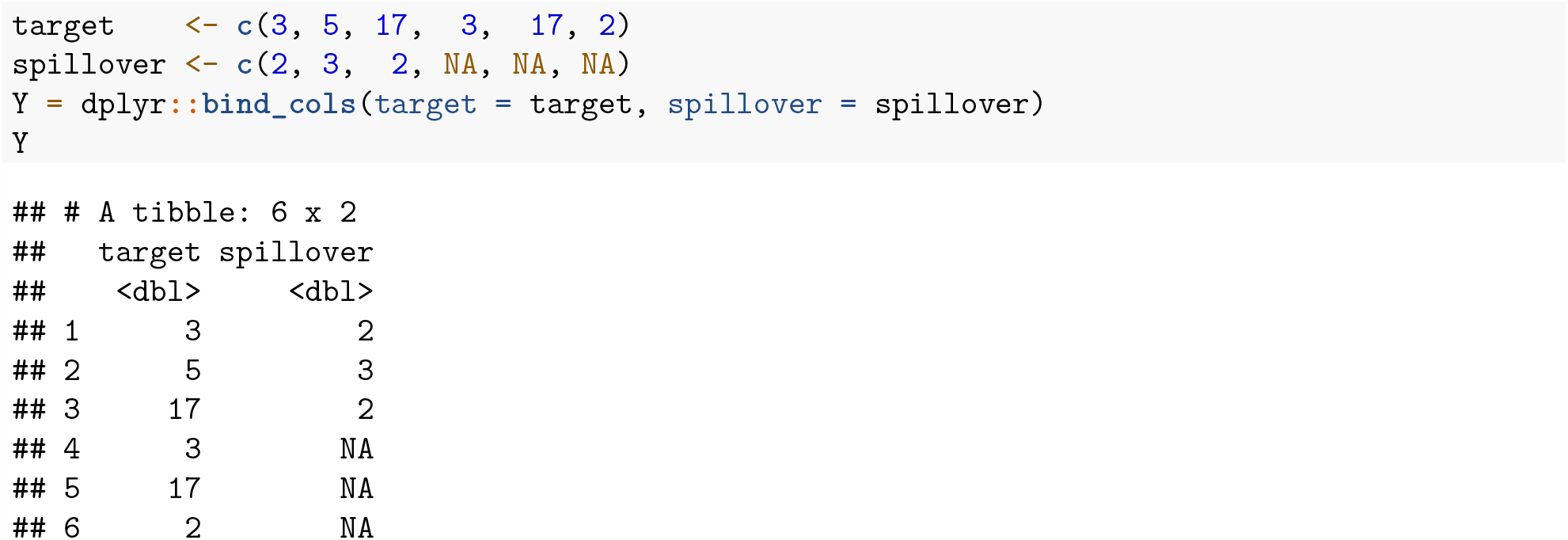

- Initialization: We initialize our EM algorithm by estimating the conditional probability of observing *y* given that it belongs to the target marker, and another conditional probability given that it belongs to the spillover marker,

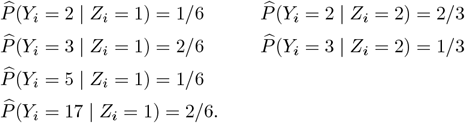

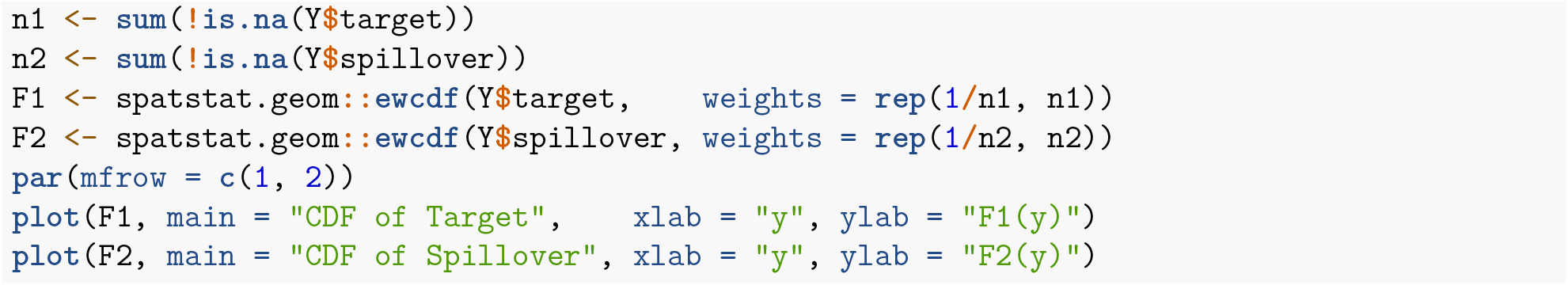

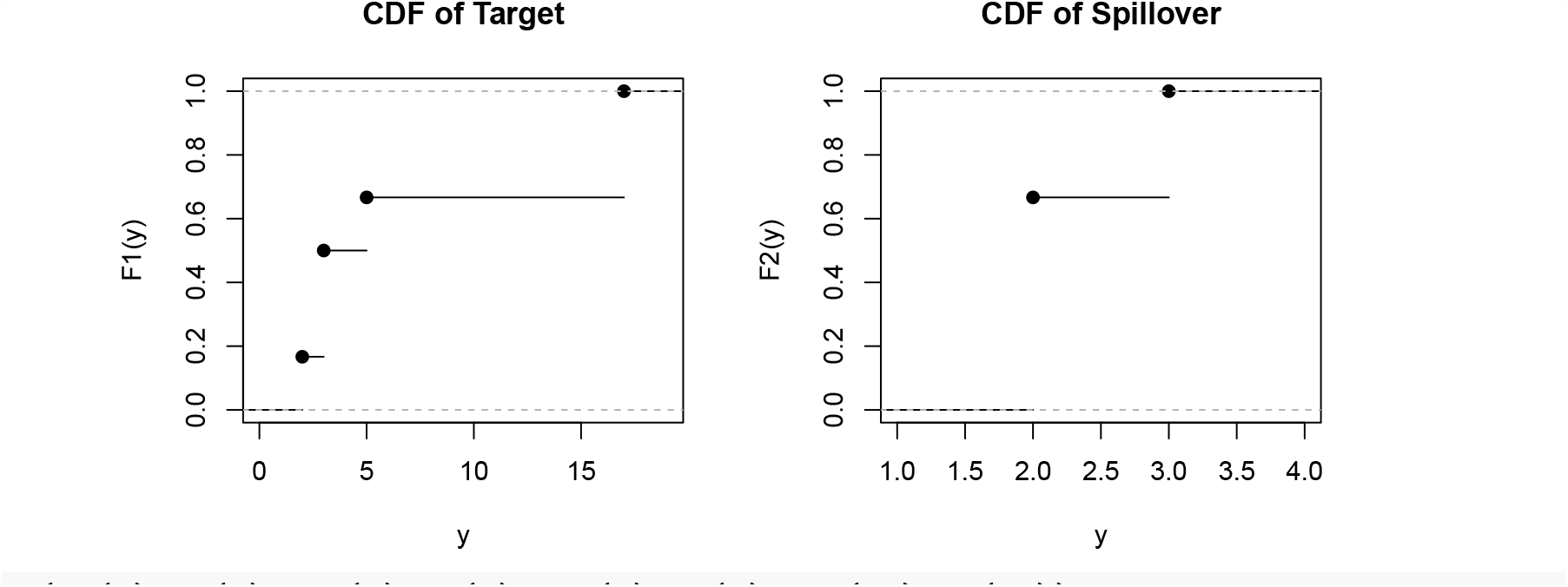

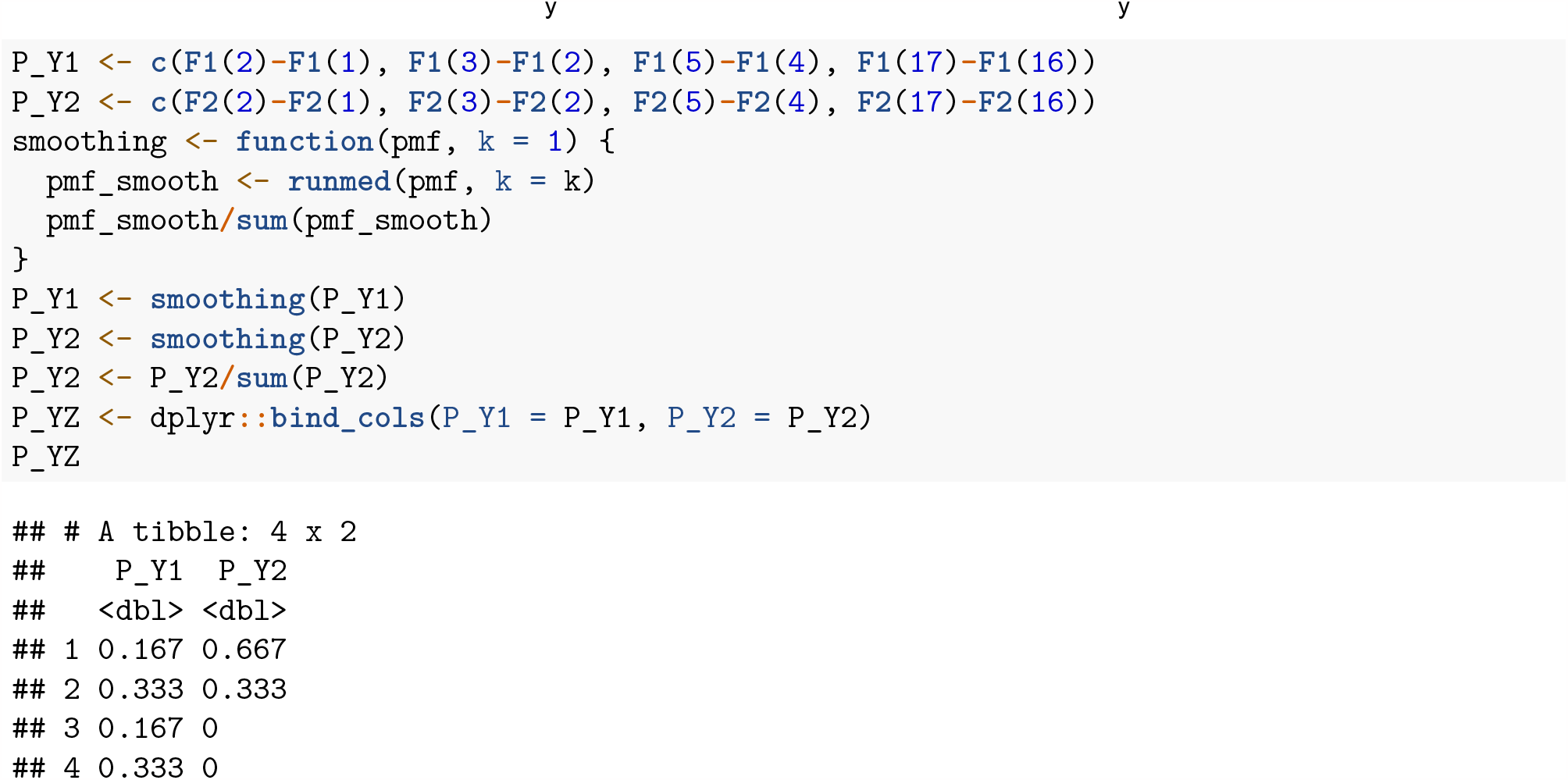

We initialize the mixture probabilities with the discrete uniform,

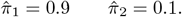

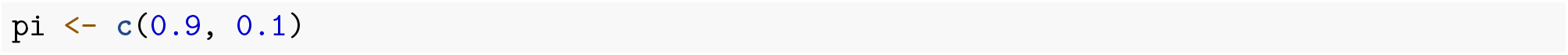 Now, we update these initial values using the E and M-steps.
- E-step: Calculate the posterior probability for the true marker, and the spillover marker,

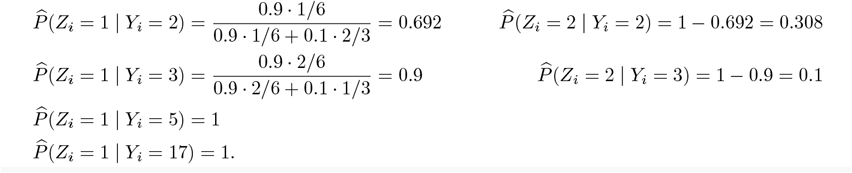

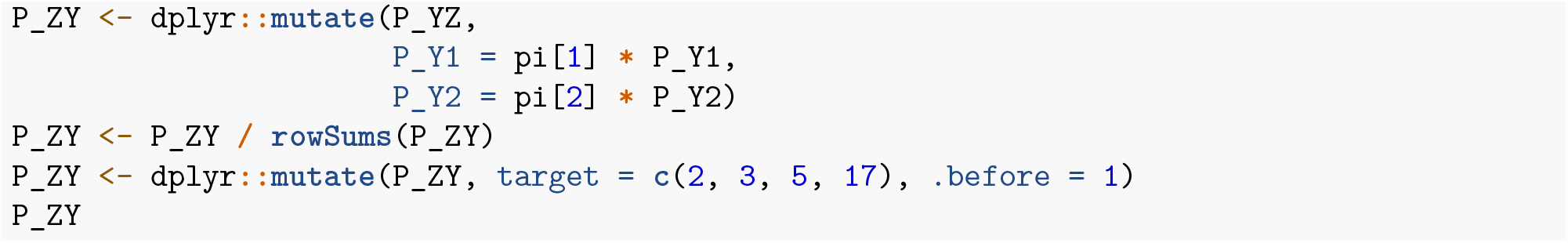

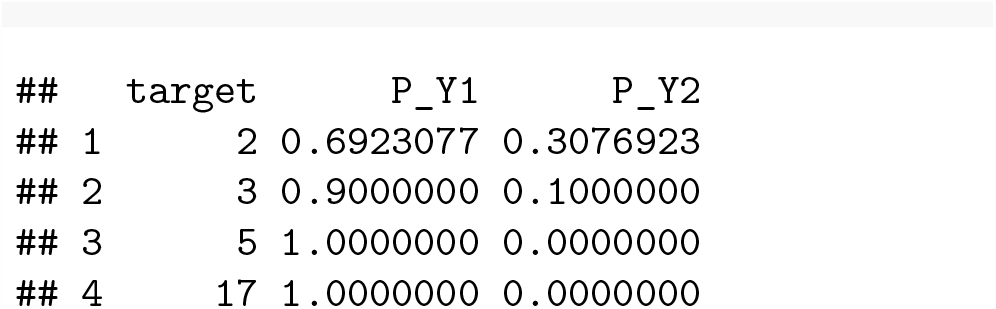
- M-step: Update the mixing probability vector,

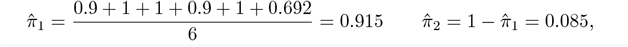

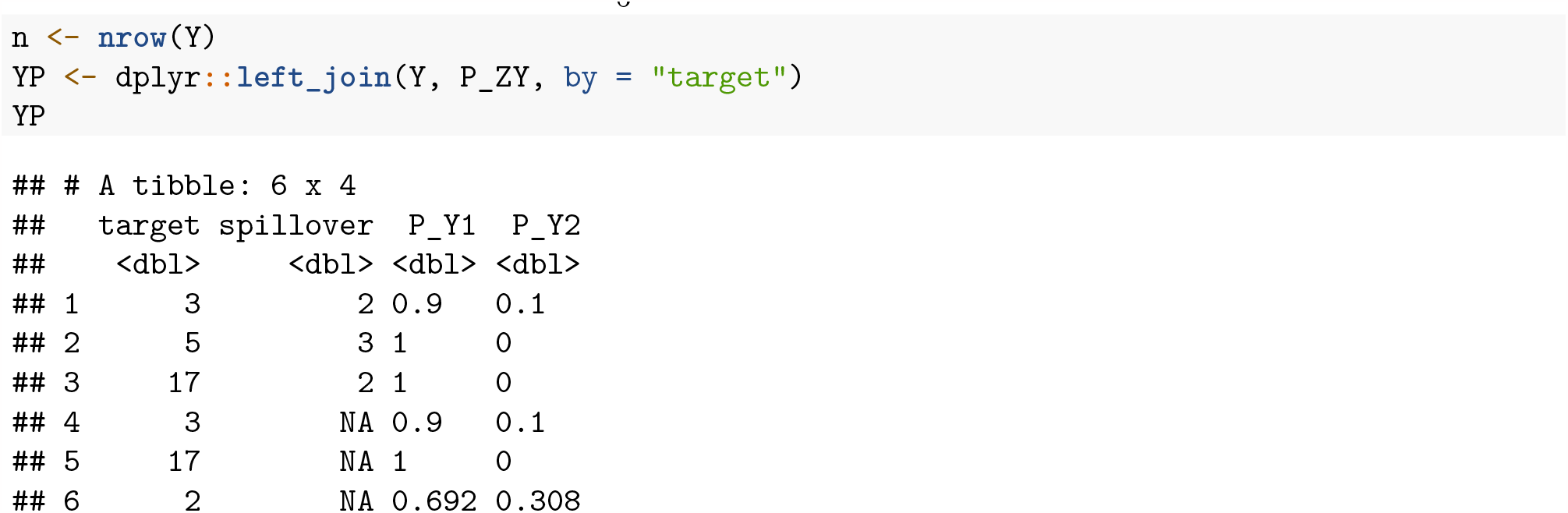

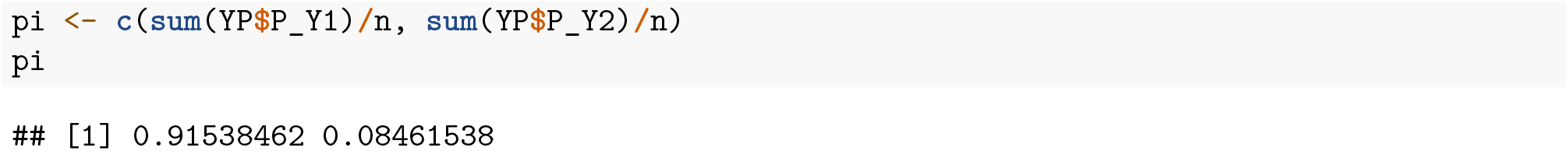

and re-estimate the distribution for the target marker using the posterior probabilities as weights, keep the non-target marker at its initial value,

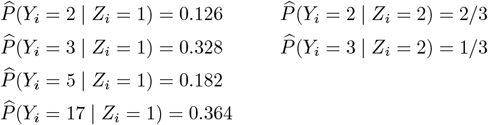

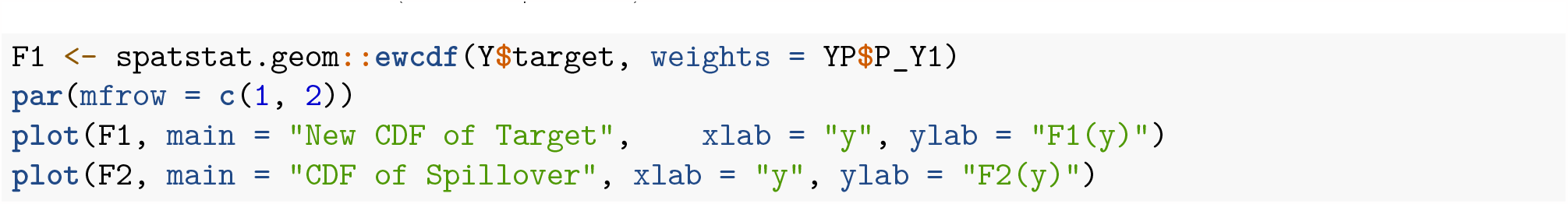

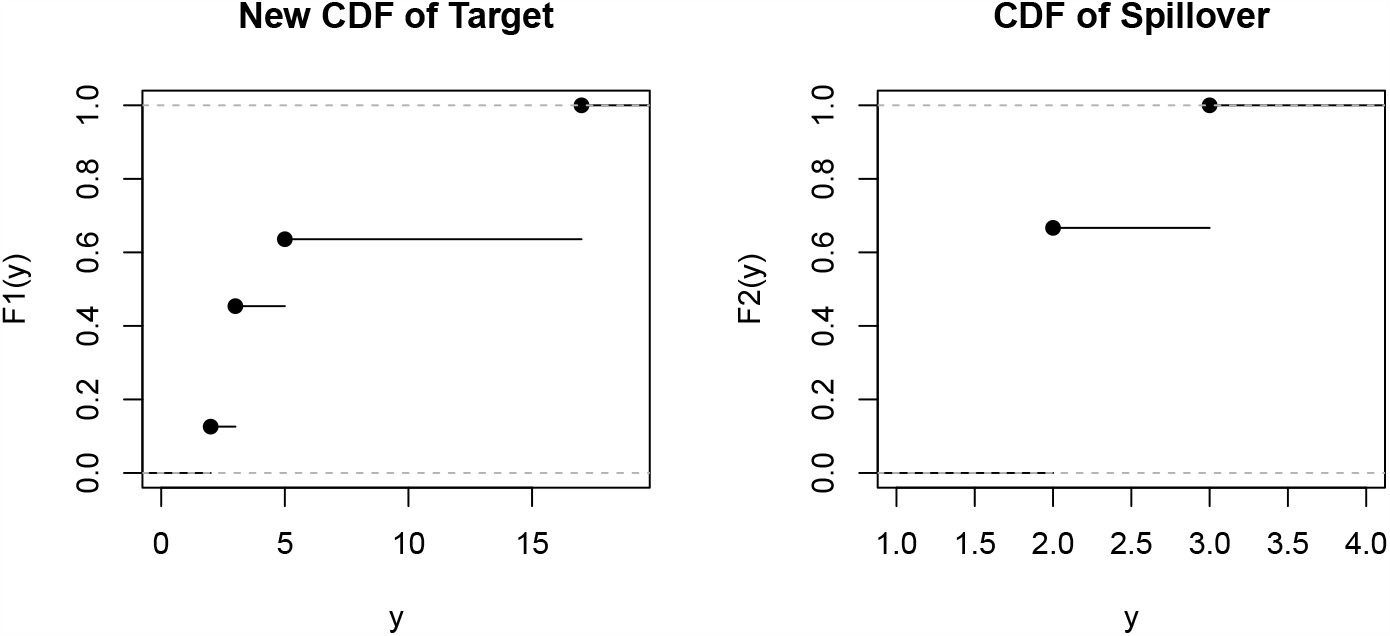

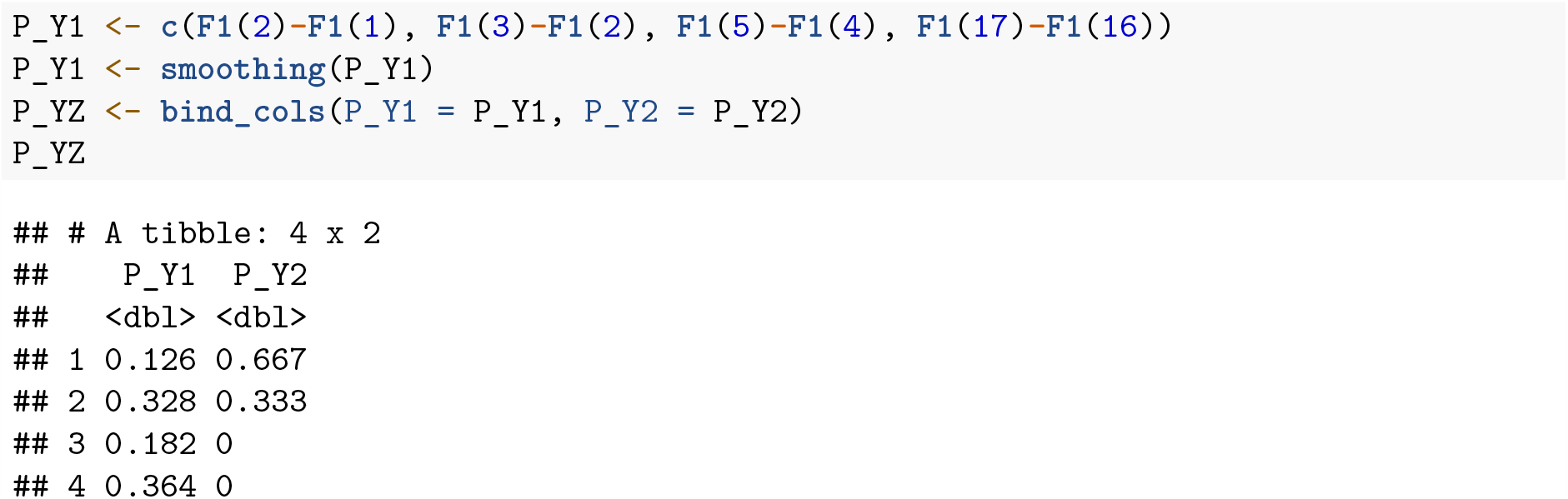

and calculate the spillover probability estimate,

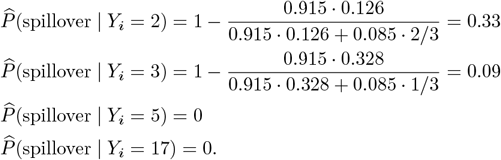

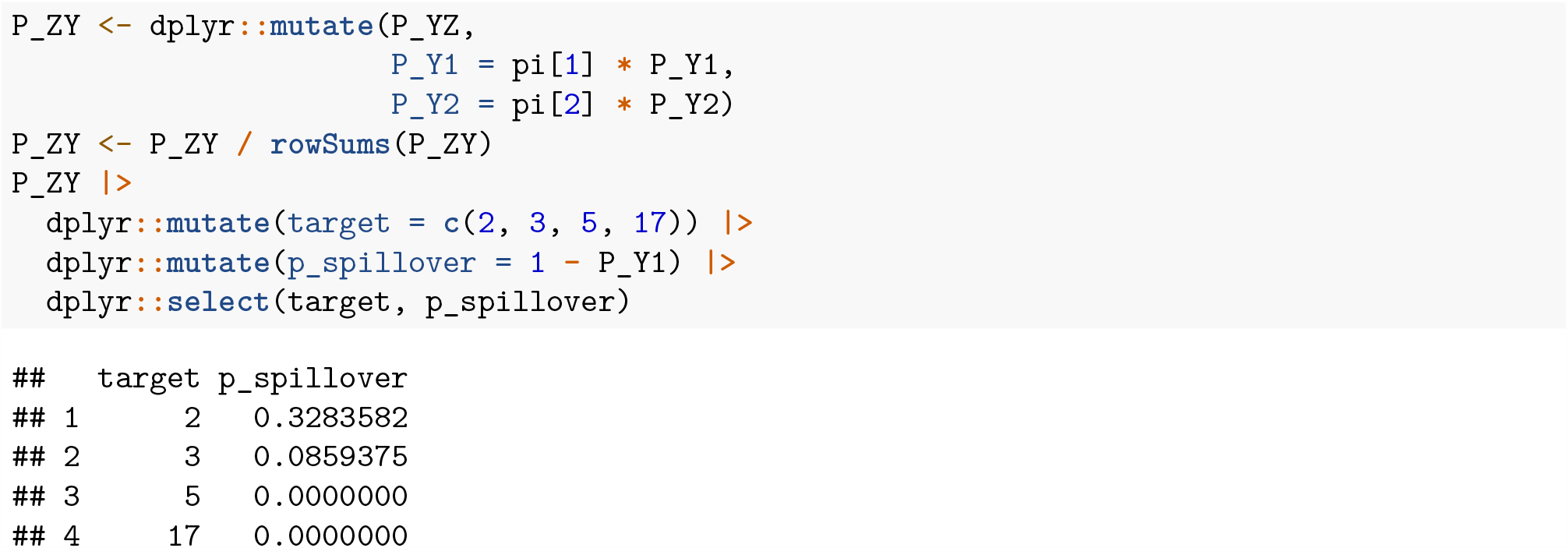

This is the result after one iteration.

## B Generative Models

### Bead Shift

Generative model for real cells *Y* of this experiment:

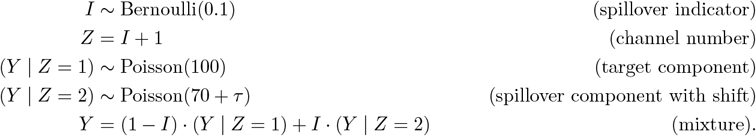

The generative model for beads is an independent copy of the unshifted *Y* | *Z* = 2 at *τ* = 0.

### Model Misspecification

Generative model for real cells *Y* of this experiment:

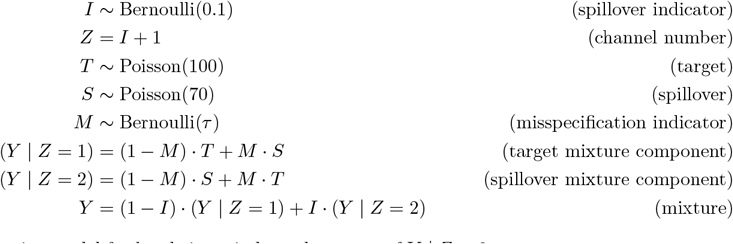

The generative model for beads is an independent copy of *Y* | *Z* = 2.

### Bimodal Spillover

Generative model for real cells *Y* of this experiment:

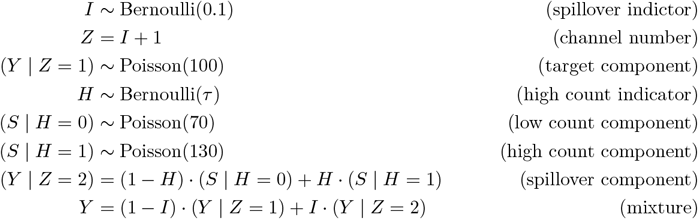

The generative model for beads is an independent copy of *Y* | *Z* = 2.

## Footnotes

1 https://github.com/marcoguazzini/spillR

2 https://github.com/ChristofSeiler/spillR_paper

## References

Bagwell, C.B. and Adams, E.G. (1993) Fluorescence spectral overlap compensation for any number of flow cytometry parameters. Annals of the New York Academy of Sciences, 677, 167–184.

Bandura, D.R. et al. (2009) Mass cytometry: Technique for real time single cell multitarget immunoassay based on inductively coupled plasma time-of-flight mass spectrometry. Analytical Chemistry, 81, 6813–6822.

Bendall, S.C. et al. (2011) Single-cell mass cytometry of differential immune and drug responses across a human hematopoietic continuum. Science, 332, 687–696.

Chevrier, S. et al. (2018) Compensation of signal spillover in suspension and imaging mass cytometry. Cell Systems, 6, 612–620.e5.

Dempster, A.P. et al. (1977) Maximum likelihood from incomplete data via the EM algorithm. Journal of the Royal Statistical Society: Series B (Methodological), 39, 1–22.

Lun, X.-K. et al. (2017) Influence of node abundance on signaling network state and dynamics analyzed by mass cytometry. Nature Biotechnology, 35, 164–172.

McCarthy, D.J. et al. (2017) Scater: Pre-processing, quality control, normalization and visualization of single-cell RNA-seq data in R. Bioinformatics, 33, 1179–1186.

McLachlan, G.J. et al. (2019) Finite mixture models. Annual Review of Statistics and Its Application, 6, 355–378.

Miao, Q. et al. (2021) Ab initio spillover compensation in mass cytometry data. Cytometry Part A, 99, 899–909.

Novo, D. et al. (2013) Generalized unmixing model for multispectral flow cytometry utilizing nonsquare compensation matrices. Cytometry Part A, 83, 508–520.

Seiler, C. et al. (2021) CytoGLMM: Conditional differential analysis for flow and mass cytometry experiments. BMC Bioinformatics, 22, 1–14.

Takahashi, C. et al. (2017) Mass cytometry panel optimization through the designed distribution of signal interference. Cytometry Part A, 91, 39–47.

Weber, L.M. et al. (2019) diffcyt: Differential discovery in high-dimensional cytometry via high-resolution clustering. Communications Biology, 2, 1–11.

